# Optic flow density modulates corner-cutting independently of age in a virtual reality steering task

**DOI:** 10.1101/2024.08.05.606519

**Authors:** Arianna P. Giguere, Krystel R. Huxlin, Duje Tadin, Brett R. Fajen, Gabriel J. Diaz

**Affiliations:** Rochester Institute of Technology, Center for Imaging Science, Rochester NY, 14620, U.S.; University of Rochester, Rochester NY, 14642, U.S.; Rensselaer Polytechnic Institute, Troy NY, 12180, U.S.

## Abstract

There is a critical need to understand how aging visual systems contribute to age-related increases in vehicle accidents. We investigated the potential contribution of age-related detriments in steering based on optic flow, a source of information known to play a role in navigation control. Seventeen younger adults (mean age: 21.1 years) and thirteen older adults (mean age: 57.3 years) performed a virtual reality steering task. The virtual environment depicted movement at 19 m/s along a winding road. Participants were tasked with maintaining a central lane position while experiencing 8 repetitions of each combination of optic flow density (low, medium, high), turn radius (35, 55, 75 meters), and turn direction (left, right), presented in random order. All participants cut corners, but did so less on turns with rotational flow from distant landmarks and without proximal optic flow. The magnitude of this flow-related effect was independent of age, although older adults cut corners more on all turns. An exploratory gaze analysis revealed no age-related differences in gaze behavior. The lack of age-related differences in steering or gaze behavior as a function of optic flow implies that processing of naturalistic optic flow stimuli when steering may be preserved with age.

## Introduction

As the number of older drivers on the road increases, so does the need to understand the effect of age on driving behavior. The average American life expectancy is 79, and more than 20% of drivers in the U.S are older than 65^1^. According to the U.S. National Highway Traffic Safety Administration, 89% of people over the age of 65 in the U.S. have a driver’s license, up from 78% in 2001^1^. Fatal crashes involving licensed drivers who are 65+ have increased from 11% to 19% from 2001 to 2021^1^. A detailed understanding of the issues that underlie these troubling statistics may present a path towards mitigation.

Although the underlying causes of age-related increases in accident rates remain unclear, changes to vision may play a large role^2,3^. Older drivers demonstrate increased rates of at-fault accidents and increased driver errors caused by a number of factors like slower reaction times, impaired visual attention, reduced visual processing speed, task switching, and decreased visual sensitivity^4–6^. There is a particularly large influence of vision on steering control^7^, and many aspects of visual perception change with age, including but not limited to changes in contrast sensitivity, color perception, and acuity^6^. Research provides convincing evidence that aging visual systems affect these functions in a way that is correlated with the number of driving accidents per mile^8,9^, and the application of an accepted assessment method suggests that this result is independent of the state of cognitive functioning. Aging visual systems are also correlated with other driving metrics like poor lane stability and increased errors in both peripheral detection and visual discrimination on the road^5,10^. These age-related changes in visual sensitivities have the potential to influence the visual control of steering.

While an age-related decline in low-level visual functions can influence steering, more complex visual functions - such as global motion processing - can also be affected by aging. Evidence suggests that aging visual systems have altered motion sensitivity^11,12^ (for a comprehensive review on visual perception and aging, see^13^). When navigating, the processing of visual motion information can play a particularly important role in the perception of heading direction^14–16^ and perceived speed^17^. Despite the critical influence of visual motion processing on steering and navigation, U.S. driver’s licenses are given based on a test of visual acuity only^6^, a perceptual dimension that has been found to only slightly impact the ability to perceive motion^18–20^. As a result, the well-documented influence of age-related changes to low-level visual functions (e.g. acuity) on driving may not be completely informative about how aging visual systems handle motion processing for steering specifically. As such, little is known about the effects of age on motion processing in a driving context.

In the present study, we tested the hypothesis that age-related changes to driving behavior can be attributed, in part, to changes in the visual processing of motion information - specifically optic flow - used to guide steering. Optic flow is a form of global motion that arises during relative movement of the observer through the visual environment. Forward movement with gaze pointed in the direction of heading results in a retinal flow field that expands radially from a point called the focus of expansion (or FoE), a term first defined by J.J. Gibson^14^. Gibson theorized that humans can navigate successfully along their desired path by visually aligning the FoE with their desired direction of heading. Although the relationships between the pattern of optic flow, perceived heading direction, and steering are far from fully understood, the decades of work motivated by Gibson’s original contributions have concretely demonstrated a role for optic flow in the visual guidance of steering^15,16,21–29^.

Prior research provides mixed evidence in support of the hypothesis that the ability to perceive heading from optic-flow degrades with age. Chou et al.^30^ measured the gait kinematics and heading direction of participants walking on a treadmill while viewing a hallway-like optic flow stimulus in virtual reality. They found that younger and older walkers responded similarly in gait and heading direction to changes in optic flow speed. Most studies, however, crafted their approach in response to evidence that the complex optic flow fields may be represented and processed as a combination of independent motion components^31,32^. These components include translational flow corresponding to left/right motion, radial motion corresponding to forward/backward movement, rotational flow corresponding to movement along a curved trajectory, and/or the added influence of gaze behavior. Some studies found a decline in motion perception with age when presented with a combination of radial flow from self-motion and rotational flow^21,33,34^ as well as translational flow^34^. Billino et al.^35^ and Atchley and Andersen^34^ discovered that responses to translational left/right motion depend on age, but the same two papers also have conflicting results regarding participant responses to radial flow fields arising from linear forward motion. They reported significant and insignificant differences respectively of age on response to radial flow. Berard et al.^36^ showed that older adults exhibit larger heading errors when tasked with walking straight in a virtual environment featuring a gradually rotating focus of expansion, suggesting that age may specifically impact behavior in response to rotational optic flow. A study by Guenot et al. suggested that motion coherence thresholds for dynamic random dot kinematogram stimuli moving left/right were similar between older and younger participants, but thresholds for radial and rotational patterns varied^37^. Together, the results of these papers point towards optic-flow-component-specific changes with age rather than an overall decrease in optic flow perception abilities.

As such, studies that provide evidence for or against changes in component-specific optic flow processing with age may have conflicting results because of differences in the stimuli of choice. For instance, in some cases optic flow was modeled using a dot field on the ground plane^21^, or a random dot kinematogram on a large screen^37^, or even by using a head-mounted display that perturbed the focus of expansion unnaturally while the participant walked straight^36^. These differences between stimuli also make it difficult to predict how they might influence behavior in a more ecologically relevant context.

Here, we chose to test the impact of age on behavior through a naturalistic steering task in virtual reality, where the quality of optic flow was systematically manipulated by changing the texture density of the virtual environment. Our choice to manipulate optic flow density was motivated by several studies that found optic flow density to impact certain functions necessary for navigation control. Warren et al.^21^ found that denser optic flow fields increase the accuracy of perceived heading direction, which suggests that density may have an influence on the utility of optic flow for the guidance of heading. One recent study used visual stimuli that varied in optic flow density to show that humans navigate towards a memorized target with greater accuracy and less variability when there is dense optic flow as opposed to sparse flow^27^. Another study focused on testing the effect of optic flow density on steering error and demonstrated that steering error increases when flow density is low^22^. Their results were consistent with the findings of^25^ that better steering performance is correlated with denser optic flow. Taken together, evidence suggests an impact of optic flow density on steering. However, flow density can vary significantly and continuously in the natural world, highlighting the need to better understand its effect on steering under more naturalistic circumstances. We predict that if age-related differences in the processing of optic flow exist, steering differences between younger and older drivers will become more apparent as optic flow density increases.

Our driving simulation required participants to steer in response to changing flow density conditions at a constant speed, so as to avoid introducing a potential confounding variable that could impact our interpretation of steering behavior in direct response to visual stimuli. In addition, our experiment leveraged virtual reality technology because VR allows for measurement of free head and eye movements as participants steer. The ability to observe human behavior without fixing the head and/or eyes generates data that better reflects natural behavior. Virtual reality also allows careful control of the presented visual stimuli, which reduces the number of confounding variables. Finally, our analysis of steering behavior was accompanied by an analysis of head and eye movements used to sample the visual environment during steering. This approach is exploratory in that it addresses the possibility of differences in steering behavior as a function of age being partially caused by differences in gaze strategies. Eye movements contribute to the pattern of optic flow incident on the retina during motion^38–40^, so it is important to consider how gaze behavior can affect heading direction judgements from flow fields and subsequently how it can affect steering performance. Our analysis will provide a first-pass level of insight into potential differences in gaze behavior.

## Methods

### Apparatus

The virtual reality simulated driving environment was developed on a computer with an AMD Ryzen 9 5950X 16-core processor, an NVIDIA GeForce RTX 3080 Ti graphics card, and 32.0 gigabytes of RAM. Data was collected at RIT using this machine. Data collection also took place at the University of Rochester using a similar machine with an 11th gen Intel(R) Core(TM) i7-11700K processor, NVIDIA GeForce RTX 3080 graphics card, and 16.0 gigabytes of RAM. Both computers ran on the Windows operating system version 10. The experiment was developed using Unity, version 2021.3.0f1. The Unity Experiment Framework (UXF^41^) was used to facilitate block and trial scheduling as well as data collection. Analysis was conducted in a Python 3.9 virtual environment using Numpy version 1.26.3, Pandas version 2.1.4, and Matplotlib version 3.8.0. The interactive environment was displayed to the user via an HTC Vive Pro virtual reality headset with integrated head tracking via Lighthouse Base station version 1. The Vive Pro has a visible field of view of 107^°^ along the horizontal, 108^°^ along the vertical^42^, with a few degrees of potential variation that accompany changes in the eyes’ position with respect to the optics and display. The Vive Pro was integrated with a Pupil Labs eye tracker set to record with a sampling rate of 120 Hz per eye and an eye image resolution of 400 × 400 pixel. Steering was controlled using a Logitech G920 steering wheel. The wheel’s SDK allowed the programmer to set parameters that influence overall sensitivity and centering spring strength, which returned the wheel to a centered orientation with adjustable resistance. Using the Logitech G Hub application, we set wheel sensitivity to 40 and centering spring strength to 40 with the intention of facilitating both comfort and control.

### Task Environment

The participant was immersed in a virtual reality task environment (Fig. 1a) in which a road, indicated by high-contrast red boundaries, followed a winding path over a flat ground plane (Fig. 1b). Although the task resembles that of driving a car, design choices were made to prioritize the potential effects of optic flow manipulations. For this reason, and insights gathered from pilot testing, a deliberate choice was made to increase exposure to optic flow in the visual field by omitting an occluding car dashboard and steering wheel from the visual environment. Similarly, the environment was rendered to be visually simple so as to reduce confounding variables that could affect steering other than optic flow.

**Figure 1.**
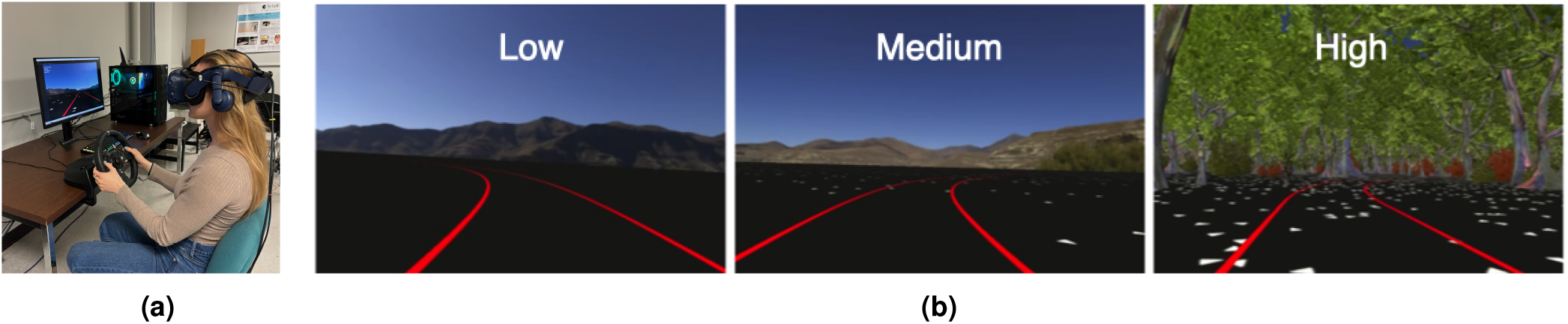
The virtual reality steering task. **1a**. A co-author steering in the virtual environment (link to video here). **1b**. View of the task environment from inside the virtual reality headset. The red lines denote the path on which participants were tasked with navigating. Each image illustrates the various levels of optic flow density, from low on the left to high on the right.

#### Road design

The road’s path was constructed by systematically piecing together individual turn segments and straight segments in an alternating pattern (Fig. 2). To facilitate trial randomization, the turns were added to the end of the road in real-time so that the road always extended two turns ahead from the road segment currently occupied by the participant. The future turns were placed at a sufficient distance to be done inconspicuously. Similarly, turns were removed from the scene immediately after the participant finished navigating them. Subjective experience and informal inquiry confirmed that this strategy of just-in-time building and removal occurred seamlessly, without participant knowledge, so the road appeared to be one continuous path. The use of a procedurally-generated roadway allowed for the systematic manipulation of independent variables between turns necessary for rigorous hypothesis testing, and it was necessary to minimize order-related effects, such as those from fatigue or task experience.

**Figure 2.**
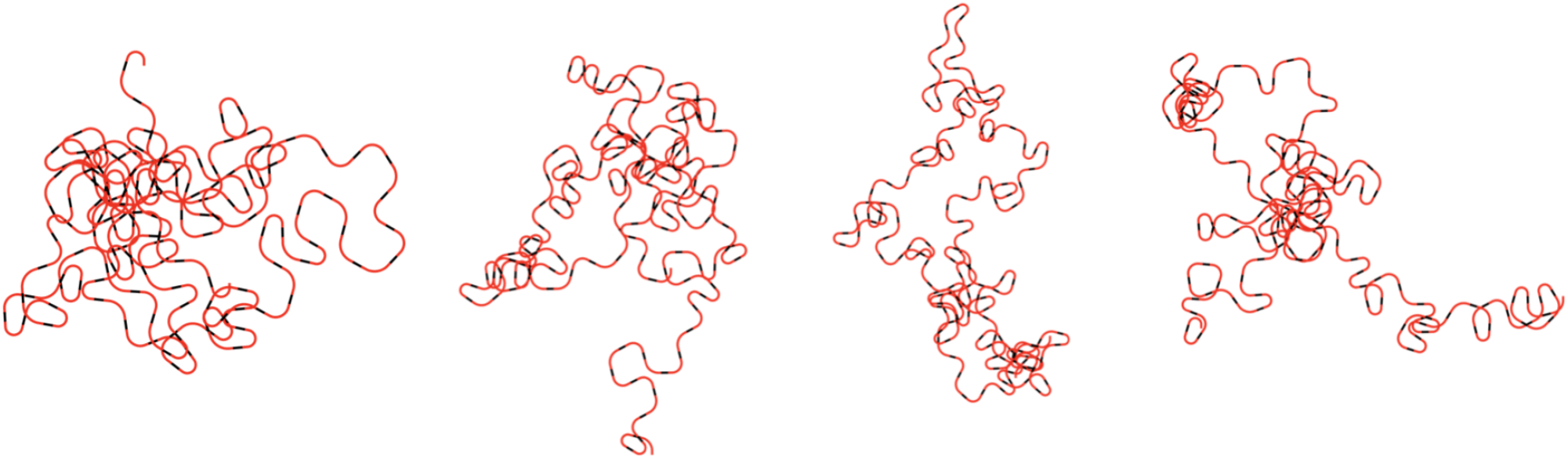
Top down views of the procedurally-generated roadway from four participants. The black segments represent the straight parts of each trial while the red parts indicate the curves, with radius of curvature 35.0m, 55.0m, or 75.0m. To prevent encounters with sections of the road that had already been traversed, the road was built and destroyed in real-time, just out of the participant’s sight.

#### Manipulation of turn direction and turn radius

Trials were equally split between left and right turns. Each road bend was 100 meters in length and defined by an arc with a radius of curvature chosen from three possible values: 35.0, 55.0, or 75.0m. Each turn was preceded and followed by straight segments of 20m in length, making each trial a total of 140 meters in length. Road curvature was varied to discourage stereotyped behavior.

#### Manipulation of optic flow density

The experimental design described below involved the manipulation of optic flow density. The density was manipulated by controlling the amount of texture present during navigation of each turn along an ordinal scale. Texture on the ground plane and the occasional presence of trees provided translational flow that could be informative about current speed and heading direction. Because the mountains were presented at a fixed distance as a part of the environmental map, the rotational flow they provided when turning was informative about turn rate, but did not yield any information about heading or the rate of self-motion.

Low optic flow density trials consisted of a black flat-shaded ground plane and distant mountains rendered via environmental map. Medium optic flow trials featured the addition of randomly placed white triangles on the ground plane, with a density of 5 triangles per square meter. On high optic flow density trials, the ground-plane triangle density was increased to 53 triangles per square meter, and a collection of trees and bushes were added to both sides of the road. In this way, we manipulated flow density by adjusting the amount of texture elements visible in the scene, and the visual changes between trial types occurred instantly between two straight-road segments of consecutive trials.

### Experiment Procedure

Each participant was instructed to “steer naturally, while centering their head and body in the middle of the single-lane road”. This instruction allowed us to measure participant lane position from several meaningful reference points including the center of the road or the inside road edge. Participants were also told to pause the experiment and/or take the headset off if they felt VR sickness.

After the experiment and task had been verbally described, each person was asked to read and sign the consent form in order to participate, and they filled out a short questionnaire about general vision, driving experience, and demographics. They were then fitted with the headset and given the opportunity to practice navigating a few turns prior to the beginning of the experiment. Approximately ten turns were sufficient for the participant to learn how to perform the task with adequate accuracy and precision.

At this point, each participant was given the option to take off the headset to re-acclimate to the natural world before beginning the experiment. An eye tracking calibration sequence and calibration assessment were completed at the beginning and end of the experiment. After the initial calibration sequence, participants could start “driving” through the environment whenever they were ready. In the event of VR sickness, the participants were instructed on how to pause themselves in between trials and how to restart when they were ready. If the headset needed to be removed, the eye-tracking calibration could be redone upon restarting.

Each turn of the virtual road encompassed various parameters, including one of three distinct optic flow conditions (low, medium, high), one of two turn directions (left, right), and one of three turn radii (35.0, 55.0, and 75.0 meters). Among these combinations, there were eight repetitions for each trial type. Participants completed 144 turns (one turn = one trial), traveling at 19.0 m/s.

### Participants

A repeated measures ANOVA power analysis using G*Power^43^ with pilot data from both groups indicated we needed 9 participants per group to produce a significant effect of optic flow on steering with a power of 0.95, alpha = 0.05. Due to participant interest and an attempt to account for VR sickness and outliers, we recruited 17 undergraduate student volunteers (mean age: 21.1 years, 14 male, two female, one nonbinary) and 13 older adults (mean age: 57.3 years, nine male, three female). Each student was eligible to participate if they had uncorrected vision or wore contacts. For our older participants, we asked that they wear contact lenses if they had them to improve performance and reduce VR sickness. Glasses were removed for the study to prevent scratching the VR lenses. Two subjects drove normally without corrected vision, two wore contact lenses for the study, and nine wore glasses. Of the eleven participants with corrected vision, ten provided their prescription information when asked, of which six were near-sighted (average left/right eye correction was -5.20D/-5.08D) and four were far-sighted (average left/right eye correction was +3.5D/+3.74D). All participants signed consent forms approved by RIT’s institutional review board and were compensated $10 for their time, unless they were participating for class credit instead. All data collection was performed in accordance with the guidelines and regulations defined in the IRB-approved consent form. Two participants were removed from the younger group due to VR sickness and four were removed from the older group for the same reason, leaving a total of 15 younger and 9 older participants with steering data to analyze.

Data from two additional participants was omitted from the gaze analysis due to eye-tracker dysfunction. Gaze data was also omitted when the average gaze validation accuracy from Pupil Labs measured over the central 16.7^°^ of the visual field was greater than 3.5^°^. This accounted for one younger adult and two older adult subjects, leaving a total of 12 younger and 5 older participants with gaze data to analyze.

### Analysis of steering behavior

To describe steering behavior, we measured steering bias as the distance from the participant’s head to the inner road edge at every time step (sampled at Unity frame rate of 90 Hz). The decision to measure position relative to the inner road edge was based on pilot data results that showed participants were cutting corners on all trials, regardless of the instruction to stay centered in the lane. Using distance from the inner road edge over time allowed us to use one metric to describe steering behavior across both turn directions. At each time step, the distance from the person’s head to the closest road vertex was recorded, and distance from inner road edge was calculated by offsetting the half-width of the road. Average trajectories could be computed from this data based on turn radius and optic flow density. Given the fact that timestamps across trials were not always perfectly sampled at equal time intervals, we interpolated across time, leveraging the “interp” function from the Python package numpy at a resolution of 90 Hz, consistent with Unity’s frame update rate. Additionally, to promote investigation of mean steering biases from inner road edge across different sections of the turns, the results were collapsed over time. We also recorded steering wheel angle and head orientation at the same sampling frequency (90 Hz), and logged all road vertices/orientation in the world for future trial-by-trial analysis.

### Analysis of gaze behavior

The Pupil Labs Core software suite was used to perform offline calibration, and to map pupil data into a gaze vector in the 3D scene. Prior to mapping gaze locations in Pupil Labs, we passed the data from each participant through EllSeg^44,45^ to increase the accuracy of 2D pupil detection. Pupil Labs maps 3D gaze by first fitting a 3D eye model of the eye to the estimated 3D eyeball centroid, and then rotating the eye-model on the basis of per-frame 2D detected pupil locations. The benefit of the relatively low-frequency updating of the 3D eye model is that the algorithm is theoretically robust to slippage. However, we found that overall accuracy was improved by first fitting and subsequently freezing the 3D model. We utilized a custom plugin to fit a 3D model to a temporal segment of the gaze data, where the specific segment varied between individuals through a process of human-in-the-loop optimization. The model was frozen after it had been fit, prior to any gaze mapping. Subsequently, gaze estimation was completed across all trials in the experiment. If a calibration sequence was present at the end of the experiment as it was for most participants, validation of gaze estimation was calculated on the final calibration sequence, otherwise it was done on the initial sequence. The average accuracy and precision of gaze estimation for each group was as follows: younger adults (n=12), accuracy=2.7^°^ precision=0.1^°^; older adults (n=5), accuracy=2.3^°^ precision=0.1^°^. After recording the validation results, the post-processed gaze data was exported into a file that included gaze location over time.

Raw gaze data was exported from the Pupil Labs software prior to being imported into a custom Python program for filtering and analysis. The data exported from Pupil Labs includes the angle of the eyes within the head for both left and right eyes, sampled at 120 Hz each. Because the cameras were not temporally synchronized, the effective sampling rate was 240 Hz, with unequal and non-constant intervals between monocular signals. Upon import, the 240 Hz Pupil Labs data was aligned with 240 Hz head orientation data through interpolation to the Pupil Labs timestamps. Samples with confidence values below 0.75 were discarded, and gaze data for that row set to *nan* (for more details on the confidence rating methodology, see Barkevich et al.^45^). Because of the presence of monocular dropouts, gaze samples could alternate between a binocular sample, monocular sample in which only one eye’s orientation was estimated at above-confidence levels, or a dropout in which a gaze estimate could not be produced at an above-confidence level in either eye. Whenever possible, monocular gaze data was converted into a single cyclopean estimate through the application of numpy’s nanmean() function across the two vectors followed by vector normalization. The cyclopean estimate of the eye’s angle within the head 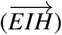 was then used to calculate the orientation of the gaze direction within world space 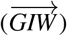 through multiplication of 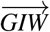, expressed in homogenous coordinates [*X*_*GIW*_, *Y*_*GIW*_, *Z*_*GIW*_, 1], with the head’s transformation matrix. To prevent translation during multiplication, the component of the head’s transformation matrix associated with translation was set to zeros. The final gaze azimuth and elevation data were prepared with a rolling mean and median filter of kernel size 9 (e.g., 37.5 ms in width) to smooth out the signal without suppressing signals of relevant saccadic behavior.

The 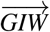 signal was converted to the spherical angles of azimuth and elevation. Gaze elevation was measured relative to the ground plane, as depicted in Fig. 3b. Azimuth was defined relative to an arbitrarily defined forward axis at which azimuth was set to zero (see Fig. 3a). This axis was sampled at the start of the experiment, after participants were instructed to hold the steering wheel and “look straight ahead”. As Fig. 4 demonstrates, gaze azimuth had a bimodal distribution consistent with the tendency for participants to look in the direction of the upcoming left or right turn. For this reason, statistical analyses of gaze azimuth magnitude was run on the absolute value of gaze azimuth.

**Figure 3.**
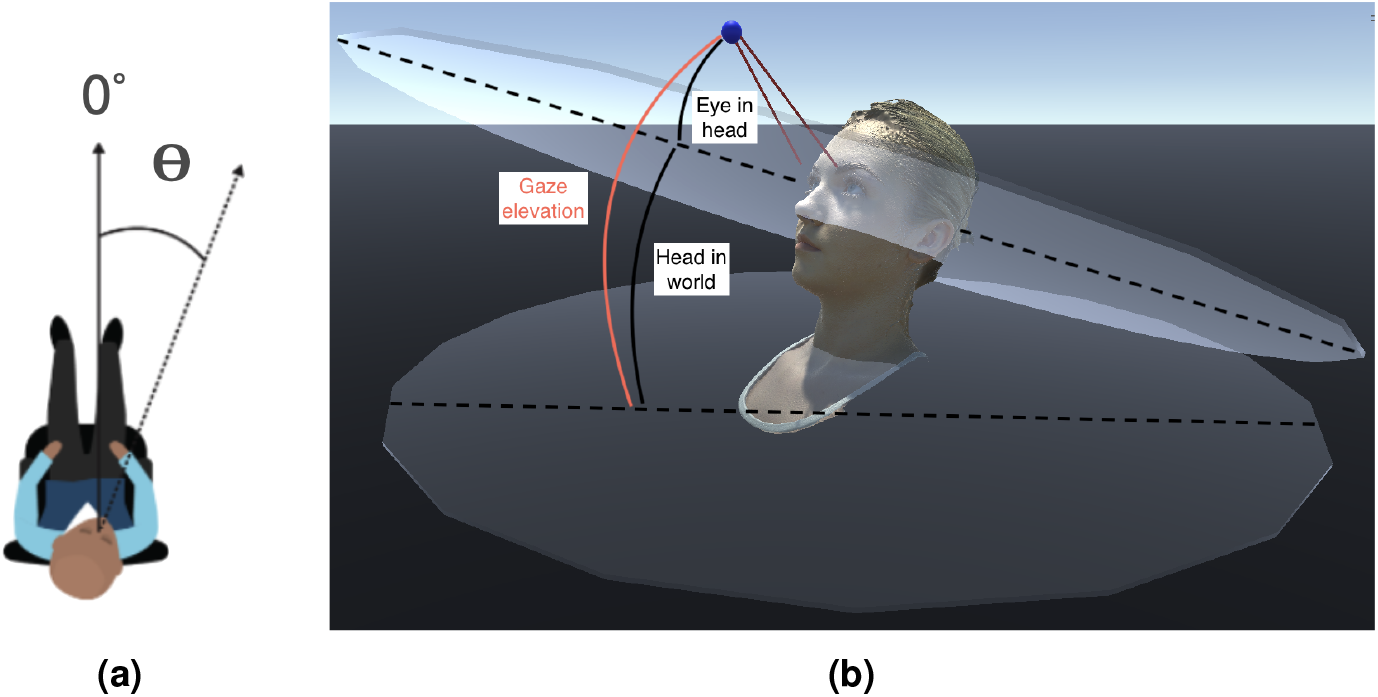
Diagrams illustrating how gaze azimuth and gaze elevation are measured. **3a**. Gaze azimuth is measured as the angle from the body forward direction (sampled upon session start) towards the left or right. **3b**. Gaze elevation is measured as the angle from the gaze point in world coordinates to the flat ground plane.

**Figure 4.**
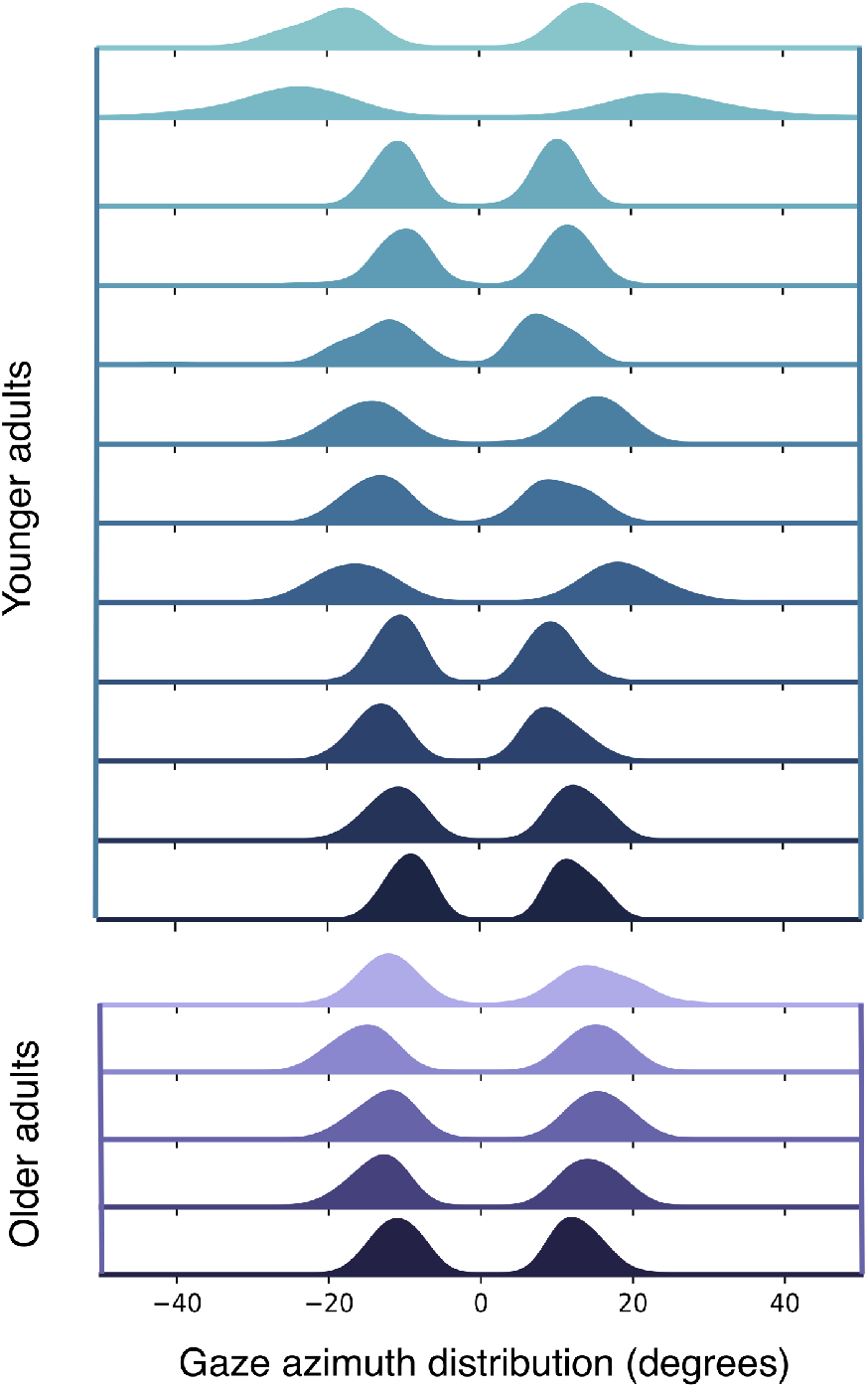
Average gaze azimuth fixations for each participant along the middle 40% of each trial. Distributions shown above demonstrate the variability of average gaze azimuth fixations across all trials for each participant which depend on turn direction (bimodal distribution) and radius of curvature of the road (statistically significant). The gaze behavior of younger adult participants is displayed on top in blue while the data from the older participants is below in purple. The figure shows variability in gaze fixation behavior across subjects.

### Statistical Analysis

A repeated measures ANOVA was run on each of the three dependent variables (steering, gaze azimuth, gaze elevation), totaling three ANOVAs. Data was analyzed across the middle 40% of each trial for all dependent variables and separated by our independent variables including turn direction, optic flow density, turn radius, and the between-group variable of subject group. In all cases, sphericity was checked using Mauchly’s test, and Greenhouse-Geisser correction was applied in the cases where the sphericity assumption was violated. The Bonferroni test for pairwise comparison of the means was used for post-hoc analyses of the ANOVA variables that reached significance. All results were evaluated for significance at an alpha of 0.05.

## Results

### Steering behavior

The left column of Fig. 5 presents representative steering behavior for individual trials from two participants. Despite the task instructions asking participants to keep their head centered in the single-lane roadway, all participants approached the inner road edge while rounding the turn, as depicted in Fig. 5. For this reason, we report road position as the *distance from inner road edge* in meters for all steering behavior results. The right column of Fig. 5 displays the gaze behavior for the same trials. This representative gaze data from two subjects captures the general tendency of participants to keep gaze elevation (green line) roughly constant throughout the turn while the azimuthal gaze angle (orange line) is shifted in the direction of the turn. The sawtooth pattern is consistent with the optokinetic nystagmus that would arise if gaze were periodically tracking features as they rotate through the visual field, similar to the behavior previously reported by^46–48^.

**Figure 5.**
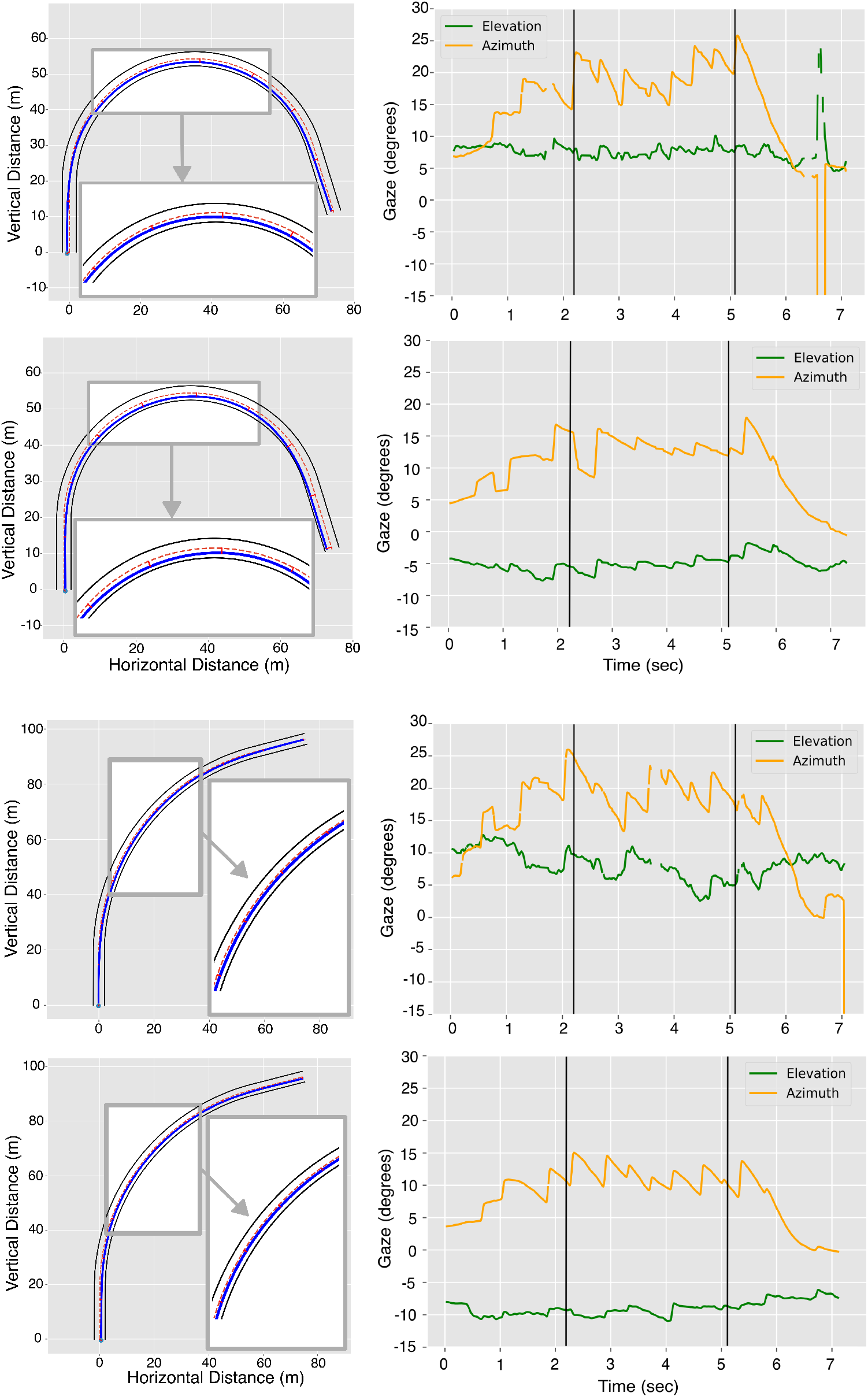
Steering and gaze data from two participants. Each row presents data from one trial. The left column depicts a top-down view of the participant’s trajectory over the course of the turn, and the right column represents the corresponding gaze behavior in degrees azimuth and elevation. All trials depict data from trials with high flow-density, but are representative of behavior that is consistent across variations in flow density. The top two rows present data from trials in which the turn radius was 35m, and the bottom two trials with a turn radius of 75m. Within each turn radius section of the figure, the top row comes from an older driver and the bottom row from a younger driver. The central 40% of the trials are indicated inside the insets of the left column and inside the black vertical lines of the right column.

The average trajectories of participants over time, separated by group and flow density, allowed for characterization of steering behavior along various segments of the turns (Fig. 6). Smaller values in this figure represent average positions closer to the inside road edge, regardless of turn direction. These representative trajectories reveal several behaviors common to all subjects. Generally, the shifts in lane position are more dramatic for sharper turns in the road, for both groups. Older participants steer with less variability than younger participants, despite having a smaller group size. In anticipation of the upcoming turns, all participants were slightly biased towards the outer road edge at time t=0, quickly crossing the center of the road at ∼0.8 seconds just before the start of the curve (at ∼1.05 seconds). Differences in steering behavior between levels of optic flow density do not begin to emerge until roughly 2-3 seconds into the turn. Distance from the inside road edge is then relatively stable until approximately six seconds into the turn. This portion of data corresponds roughly to the duration for which the participant was negotiating the most central part of the bend in the road. For this reason, the subsequent analysis of driver steering performance and the accompanying gaze behavior is applied to the central 40% of the trial. Because speed and turn length were held constant, the central 40% corresponded to approximately 2.2 to 5.1 seconds of the turn duration across all conditions. This portion of the trajectory is visually indicated in Fig. 6 as black horizontal lines and in Fig. 5 by magnified insets.

**Figure 6.**
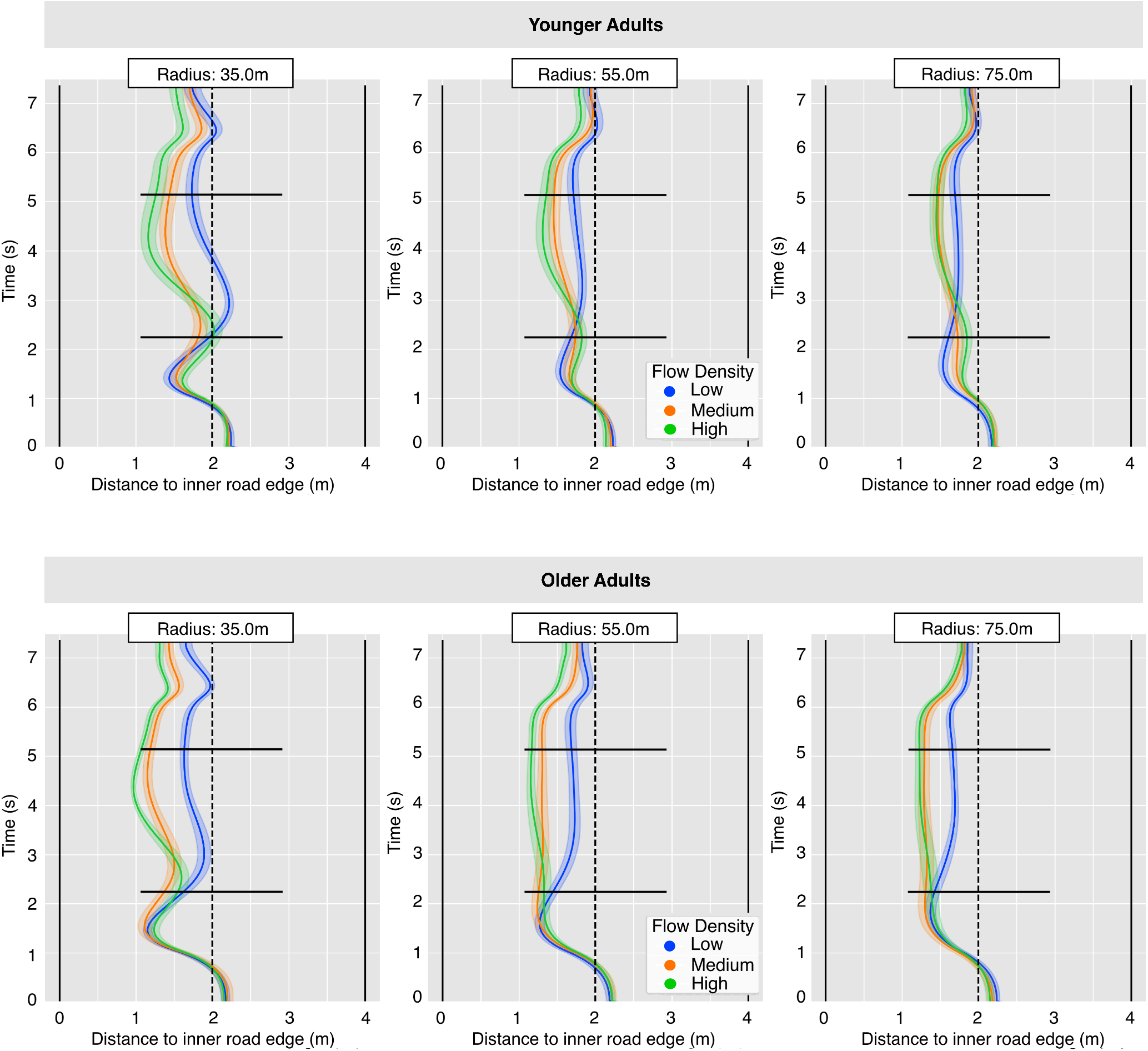
Average steering bias over time, as measured from the inner lane for younger drivers (top row, n=15) and older adults (bottom row, n=9). Shaded areas represent 95% confidence intervals. On the ordinate axis, t=0 represents the trial start. The curve is roughly along t=1 to t=6 seconds with straight parts leading into and out of the turns from t=0 to t=1, and t=6 to t=7. Black vertical lines represent the edges of the 4m single-lane roadway. The center of the road is marked by the vertical dotted line, although no such indication was presented in the simulation. Black horizontal lines indicate the beginning and end of the central 40% of the turn.

All participants showed a general tendency to cut towards the inside road edge (e.g. to corner-cut) while rounding turns, and they did so by a greater margin on left turns (Fig. 7a), as indicated by a significant effect of turn direction (F(1, 22) = 21.7, p *<* 0.001,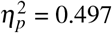). Older adults consistently engaged in more corner-cutting than younger adults, irrespective of turn direction (see Fig. 7a). This was reflected by smaller average distances from the inner road edge (effect of subject group on average lane position: F(1, 22) = 7.63, p = 0.011,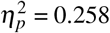).

Although statistical analysis with a mixed-design ANOVA revealed a main effect of optic flow (F(1.382, 30.404) = 70.82, p *<* 0.001,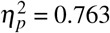), the interaction of age group and optic flow failed to reach significance (F(1.382, 30.404) = 2.42, p *<* 0.121,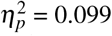). The relevant data are presented in Fig. 7b. Post hoc tests of flow condition indicate that differences lay between the low flow condition and both the medium and high levels of optic flow (low/medium: p *<* 0.001, low/high: p *<*, medium/high: p=0.333).

**Figure 7.**
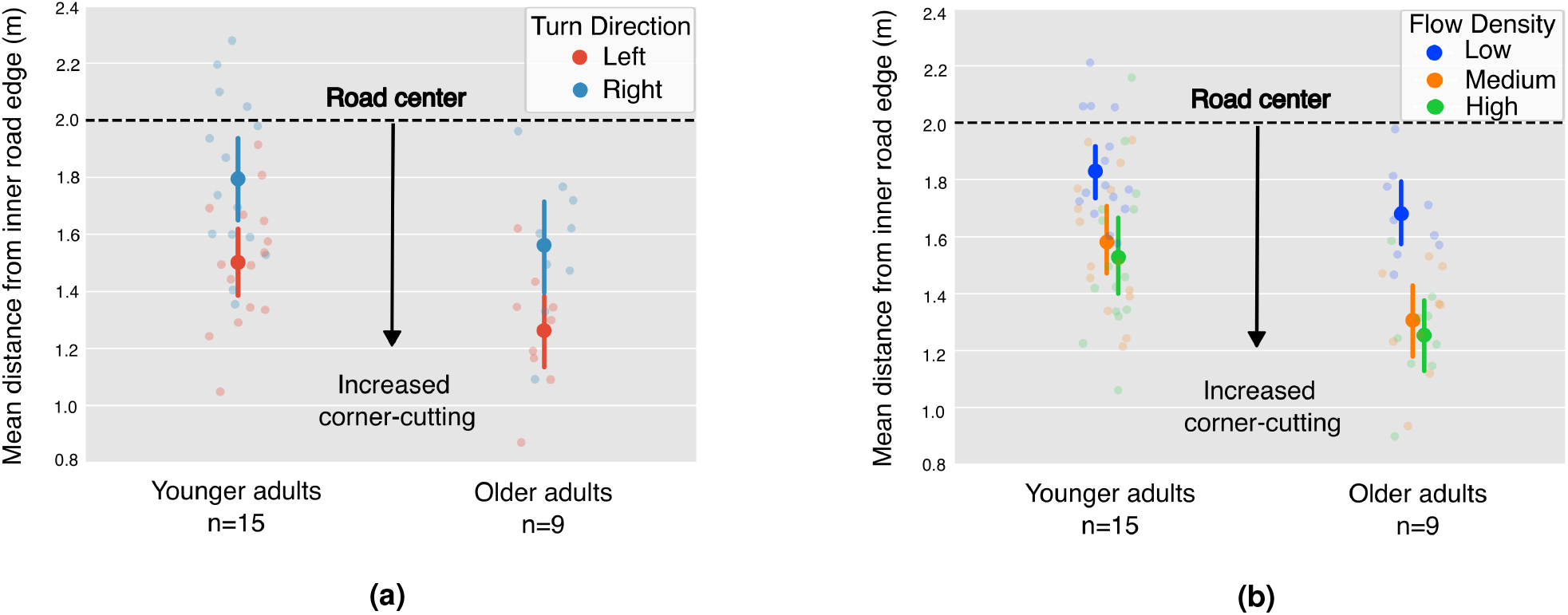
Average distance from inner road edge for each group. Averages are computed over the middle 40% of each trial. Error bars represent the 95% confidence interval of variability across subjects within each group. **7a**. Data separated by group and turn direction, but averaged across optic flow **7b**. Data separated by group and level of optic flow density, but averaged across turn directions.

There was also a significant effect of turn radius on average lane position, but only through an interaction with optic flow density (interaction of turn radius and optic flow: F(3.155, 69.402) = 16.83, p *<* 0.001,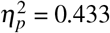). Post hoc results revealed that steering behavior differs between the sharpest turn radius and the other turn radii only under the low flow density condition (low flow: 35m/55m: p = 0.004, 35m/75m: p *<* 0.001). This result suggests that the sharpest turn radius impacts the amount of corner-cutting relative to the other radii only when flow from self-motion is absent (low density condition), otherwise it does not impact steering performance.

### Gaze behavior

Because driver steering behavior was analyzed across the middle 40% of all turns, we analyzed gaze behavior over the same segment of each trial. Eye movements can affect retinal flow^39^ which has the potential to affect steering, so analyzing across the same segment is necessary. From the right column of Fig. 5, it is evident that participants exhibited signs of nystagmus within the central 40% of each trial. Despite the nystagmus patterns, averaging gaze location within the central 40% is justified because we are interested in where people steer and look on *average* while they navigate the central part of each turn.

Statistical analyses of average gaze azimuth were conducted on the absolute value of fixation location (i.e., on its magnitude), as discussed in the gaze analysis subsection of the methods. The effect of turn radius on gaze azimuth is significant across both groups (F(1.10, 16.56) = 182.39, p *<* 0.001,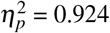). Fig. 8a illustrates this effect, where subjects in both groups generally looked at more eccentric locations when navigating sharper turns. Neither turn direction nor optic flow (Fig. 8b) affected gaze azimuth. The interaction of optic flow and subject group was significant (F(1.89, 28.40) = 5.58, p = 0.010,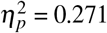), but post hoc analysis showed no significant pairwise effects with group and flow density conditions. Lastly, an analysis of the gaze elevation data for all participants revealed no significant effects between groups or any other independent variables.

**Figure 8.**
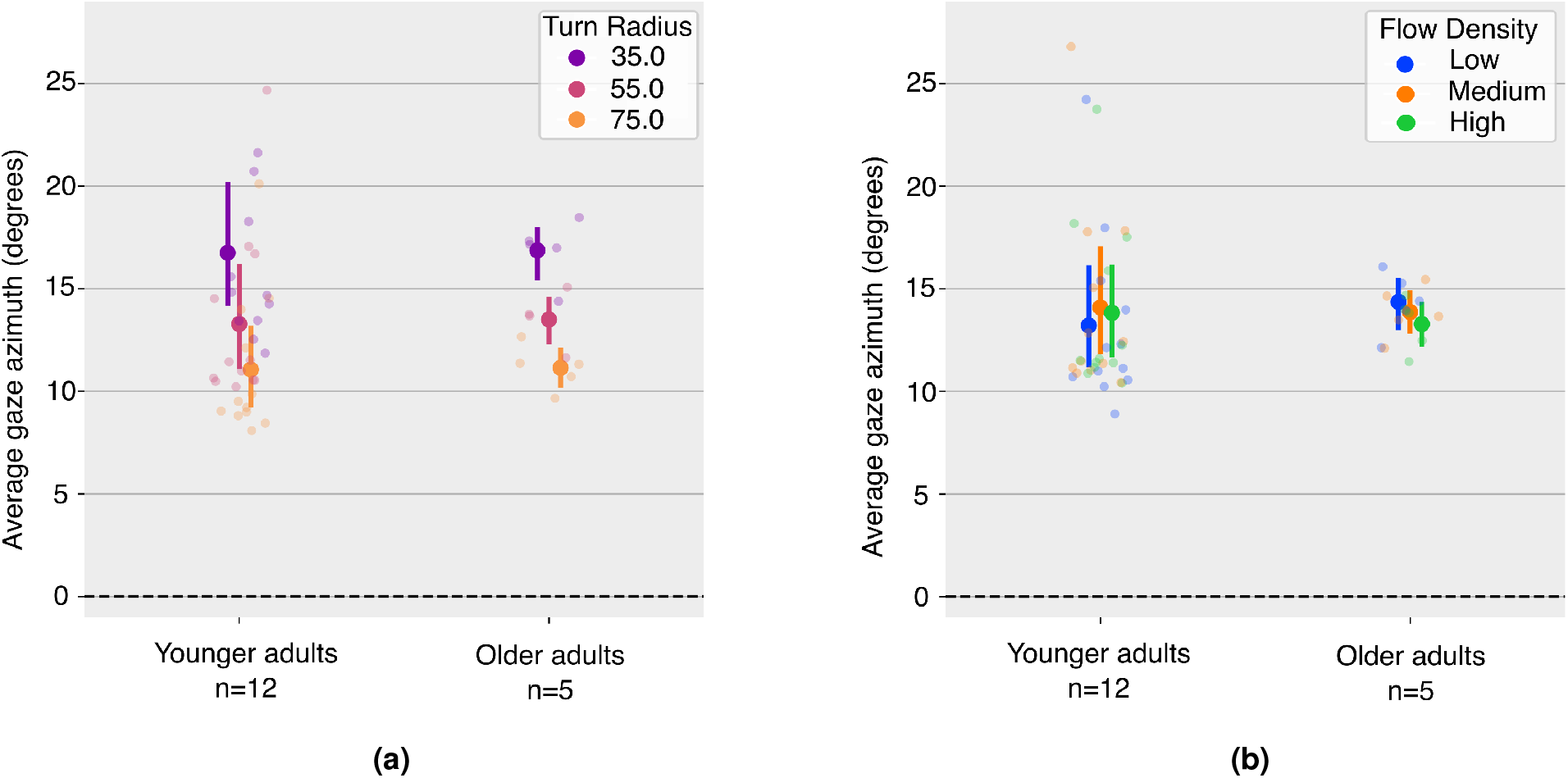
Average gaze azimuth locations over the middle 40% of each trial only. **8a**. Data is separated by turn radius. There is a significant effect of turn radius on gaze azimuth. Participants look more into the periphery on sharper turns, and this is consistent across both groups. **8b**. Data is separated by optic flow. There is no significant effect of optic flow on gaze eccentricity. Error bars show the 95% confidence intervals of variability across the participants in each group.

### The coupling of gaze to steering

While the results presented above focus on analyzing steering and gaze separately, the purpose of this subsection is to briefly illustrate the relationship between gaze and steering, given that the two variables may be coupled^49,50^. Fig. 9 presents the relationship of gaze azimuth to wheel angle for all subjects. Although a correlation exists between gaze and wheel angle by turn radius (Pearson’s correlation: r(49) = 0.556, p*<*0.001), there is no evidence of a correlation within turn radius (Pearson’s correlation: r(49) = 0.118, p=0.416). This suggests that, although gaze azimuth varied with the coarse changes in steering behavior that are required by categorical changes in turn radius, there was no evidence of fine-grained, trial-to-trial coupling of gaze azimuth and steering behavior within a single turn radius.

**Figure 9.**
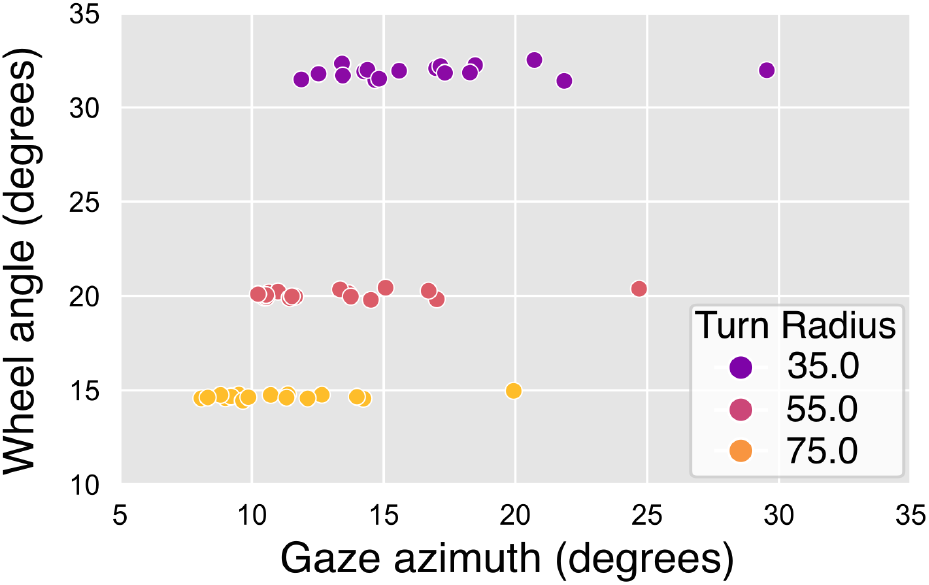
Wheel angle and gaze azimuth are tightly correlated, suggesting that participants look where they want to steer. Wheel angles are greater to successfully navigate tighter turns. This data is consistent with the point-attractor theory of Wilkie and Wann^50^. However, the data within each turn radius are uncorrelated, suggesting that the point-attractor theory may only apply on a coarse scale. One participant looked noticeably further in azimuth relative to others, but their gaze behavior patterns are otherwise consistent with the rest.

## Discussion

This study tested the hypothesis that there are age-related differences in the processing of optic flow for the visual guidance of steering. We found that younger and older drivers responded similarly to systematic variations in the density of texture elements that form the optic flow field during translation when driving. Our findings build upon a long history of research that has produced mixed results about the role of age on the processing of optic flow. Our main result, that age does not significantly impact steering behavior as a function of optic flow density, is consistent with prior results that used more conventional psychophysical methods to identify motion discrimination thresholds^37^ and psychometric thresholds for FOE location judgments^35^. Although this study focused on steering behavior, our results are consistent with the finding that responses to optic flow stimuli while walking in VR are similar for both younger and older participants^30^, and with prior results that show no differences in walking speed for older adults in response to changes in optic flow speed^51,52^. However, other studies report the conflicting finding that accuracy of heading judgments from optic flow decreases with age^21,33^ and motion discrimination thresholds increase^34^. These differences may be attributed to differences in the visual stimuli used. For example, the stimulus used by Lich and Bremmer^33^ was composed only of 3D dot fields in VR, in the absence of a ground plane or other environmental structure. Similarly, the studies by Atchley and Andersen^34^ and Warren et al.^21^ involved stimuli displayed on conventional 2D desktop monitors, either as random dot kinemateograms or as white pixels on a blue ground plane.

Rather than attempt to resolve the inconsistencies about age-related optic flow processing in previous work, our study attempted to investigate age-related optic flow processing in the context of steering, a task we chose for its ecological relevance to the real-world task of driving. We created a 3D environment with simulated objects from the natural world, rendered to scale, and presented stereoscopically on a Vive Pro VR headset with a wide field of view. Peripheral vision beyond *>* 50^°^ is also sensitive to motion information^53,54^ and has been shown to contribute to optic flow perception^55–57^. Optic flow was structured by the 3D environment and, similar to natural contexts, varied in texture density.

The use of eye tracking in VR allowed us to test the possibility that age-related detriments to optic flow processing exist, but are diminished in older adults through the use of compensatory gaze behavior. This is possible because eye movements contribute to the pattern of motion that forms the optic flow field, and opens the possibility that intelligent gaze behavior might actively bring about task-relevant patterns of flow that are, for example, informative of future heading^39,58^. However, our exploratory analysis failed to find differences in gaze behavior between groups, and thus offers no support for this hypothesis. It should be noted that due to technical issues related to eye tracking in VR, group sizes were unequal and data from only five older drivers contributed to the final analysis of gaze behavior. This poses a potential challenge to statistical power. Regardless, the gaze data collected and analyzed showed remarkably little variation across conditions.

Although the data do not support the interpretation that optic flow processing degrades with age, it does motivate a novel hypothesis for how age might lead to more accidents in the natural context. We found that all drivers hugged the inside road edge, with older adults cutting corners to a greater extent. We also found additive effects of other variables on lane position biases, resulting in the observation that corner-cutting is greatest for turns to the left as well as for turns with medium or high optic flow density. Our environment featured a single-lane road, but corner cutting on double-lane roads might instead result in a pattern of behavior where older drivers hug the inside road edge and, on leftward turns, position themselves closer to a lane of oncoming traffic, possibly leading to a higher rate of collisions.

Given that the results of this study do not support the theory that there are age-related differences in the way people utilize optic flow to steer and modulate corner-cutting, we offer several alternative explanations for why older adults might engage in more exaggerated corner-cutting. One possibility is that corner cutting is a response to VR sickness and a desire to shorten the path traveled. This seems unlikely, given that speed was constant, and the reduction in experiment time would have been negligible. Moreover, we observed that some individuals who did not experience any VR sickness for the entirety of the experiment also exhibited consistent corner cutting behavior. However, because the relationship between susceptibility to simulator sickness and driving behavior was not directly tested in this study, this counter-argument remains purely anecdotal. One final possibility, and the one that we find to be most plausible, is that corner-cutting is simply a learned behavior that increases in magnitude with driving experience. Many previous studies provide evidence that humans cut corners when steering^59–61^. The fact that participants cut corners in this study despite task instructions asking them to stay centered in the lane implies that corner cutting could be a more natural steering tendency. The significant difference that subjects displayed in the amount of corner cutting on left vs. right turns also has implications for real-world driving. We speculate that the tendency to cut corners more on left turns exists because all our participants currently drive in the U.S. and are used to positioning themselves in a driver’s seat that is on the left side of the car. A follow-up study involving drivers from countries with driver’s seats on the right could test this hypothesis.

Our data also revealed that all participants, regardless of their age, cut corners to a greater extent on trials with medium and high optic flow density compared to the low optic flow density trials. A potential reason for this behavior is that the reduction of optic flow density in the surrounding environment provides a sensation of slower perceived speed^62^ which could influence lane position. Indeed, a prior study^63^ directly manipulated the magnitude of optic flow as drivers navigated turns and found that perceived flow speed affects average driver position in a lane. It is worth noting, however, that the study did not directly manipulate global flow speed uniformly, but instead introduced flow asymmetry on one side of the road while drivers negotiated the curves. The authors found that average lane position did not depend on the asymmetric patterns of flow, but instead it depended on the global average of flow speed, which positively correlated to amount of corner-cutting^63^. Although the present study manipulated flow density instead of flow speed, Snowden et al. found that increasing flow density also increases perceived speed^62^. Together, these observations suggest that our manipulation of flow density may have affected perceived speed, causing drivers to engage in greater corner cutting which is consistent with the results of^63^. Anecdotally, several participants commented, without being prompted, about the sensation of traveling faster during the medium/high flow conditions. The feeling of changes in perceived speed also aligned with the subjective experience of the authors. To strengthen this interpretation, we conducted a post-hoc analysis on changes in wheel kinematics in response to changes in perceived speed for each turn radius and flow density condition, and the results revealed that participants turned the wheel more slowly when flow density was lower (see Supplemental Fig. S1), as though an underestimation in speed reduced the urgency of bringing about the desired change in heading. An alternative explanation for why participants cut corners more on medium/high optic flow density trials could be that the absence of flow from self-motion caused a qualitative shift in the visuo-motor strategy used to drive. This effect was previously observed in^64^ which also found that humans rely less on optic flow to navigate as the signal became more sparse or unreliable.

Although it was not the central aim of this study, the data provide some insight into the role of gaze in the visual guidance of steering. Perhaps the most developed theory that remains largely supported by empirical data proposes that gaze serves as a point-attractor for the future desired trajectory^50^. Simply stated, the theory predicts that drivers steer where they look, so changes in steering behavior should be accompanied by similar changes in gaze direction. To some degree, our data support this pattern. The magnitude of gaze azimuth is positively correlated with turn radius (Fig. 9), which is to be expected if participants are simply looking a fixed distance or duration ahead. However, the theory is inconsistent with the observation that within each turn radius, gaze angle showed no trial-to-trial covariation with wheel angle. The point-attractor theory is also inconsistent with the finding that steering behavior is affected by the absence of translational flow from self-motion (i.e., in the low flow condition) but gaze behavior is not. It is possible that the point-attractor theory applies coarsely in the case of changing road curvature but is independent of optic flow density. This interpretation is strengthened by the fact that the correlation between gaze and steering within each turn radii is also insignificant for only the trials including translational flow from self-motion (*r*(49) = 0.138, *p* = 0.325).

Overall, our results support the interpretation that age does not impact steering or gaze behavior as a function of optic flow density in this task. Our primary analysis of steering data provides convincing evidence that aging visual systems do not impact the processing of optic flow required to control steering in our virtual environment. That said, age does seem to be positively correlated with the amount of corner cutting when steering, and this is likely a result of increased driving experience that comes with age. We detected no other significant, age-related differences in steering or gaze behavior.

## Data Availability

Access to compressed pre-processed data that includes all behavioral metrics described in this manuscript can be found on GitHub [here](https://github.com/ari-g-rit/Public-Data----The-effect-of-optic-flow-density-on-steering-in-VR-by-age.git). The repository includes one data file for each participant as well as the analysis file that can be used to read in the data and visualize results. Raw data is available from the corresponding author upon request. A video of the task can be found [here](https://www.youtube.com/watch?v=_3Mginft_zk).

## Acknowledgements

This research was supported by the Research to Prevent Blindness/Lions Clubs International Foundation Low Vision Research Award (LVRA). We would like to thank Chrys Callan at the University of Rochester for her regulatory support.

## Author contributions statement

All authors designed the experiment, and A.G. conducted the experiment. A.G and G.D. analyzed the data. A.G. and G.D. prepared the manuscript, and all authors reviewed and edited the manuscript.

