## Supplemental Fig. S1 for "Optic flow density modulates corner-cutting independently of age in a virtual reality steering task"

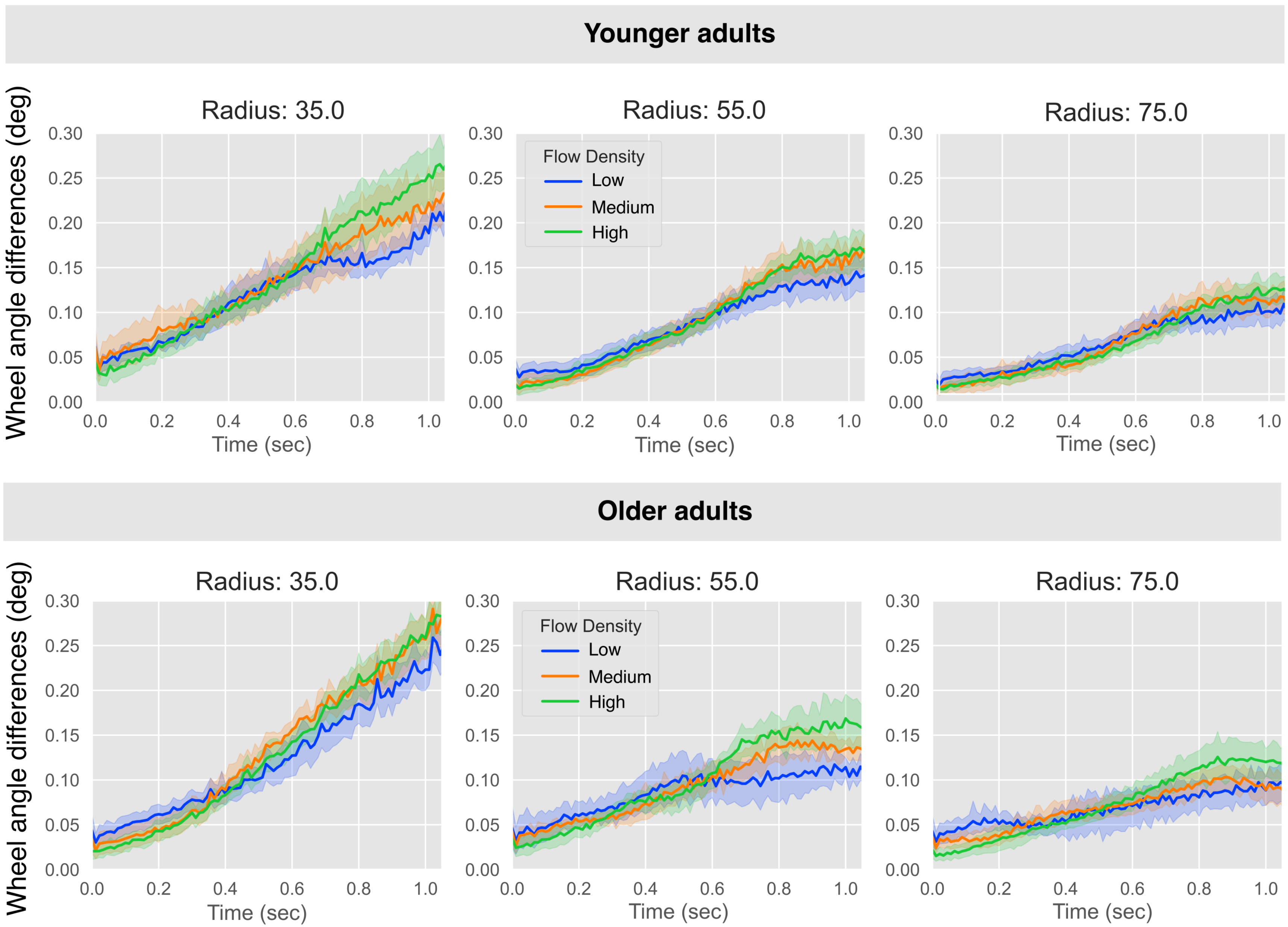


**Figure S1:** Sample-to-sample differences in wheel angle over time. The figure displays the beginning of each trial during which participants crossed over the 20m of straight road segment that preceded the turn. Data are separated by group, turn radius, and optic flow. Differences in turning rate begin to emerge around 0.6 seconds, approximately 0.4 seconds prior to entering the turn. The observation that turning rate was lowest for the low-flow condition suggests that subjects may have underestimated their speed, thus reducing the urgency to turn the wheel for successful navigation of the upcoming turn.
